# Lightweight taxonomic profiling of long-read metagenomic datasets with Lemur and Magnet

**DOI:** 10.1101/2024.06.01.596961

**Authors:** Nicolae Sapoval, Yunxi Liu, Kristen D. Curry, Bryce Kille, Wenyu Huang, Natalie Kokroko, Michael G. Nute, Alona Tyshaieva, Alexander Dilthey, Erin K. Molloy, Todd J. Treangen

## Abstract

The advent of long-read sequencing of microbiomes necessitates the development of new taxonomic profilers tailored to long-read shotgun metagenomic datasets. Here, we introduce Lemur and Magnet, a pair of tools optimized for lightweight and accurate taxonomic profiling for long-read shotgun metagenomic datasets. Lemur is a marker-gene-based method that leverages an EM algorithm to reduce false positive calls while preserving true positives; Magnet is a whole-genome read-mapping-based method that provides detailed presence and absence calls for bacterial genomes. We demonstrate that Lemur and Magnet can run in minutes to hours on a laptop with 32 GB of RAM, even for large inputs, a crucial feature given the portability of long-read sequencing machines. Furthermore, the marker gene database used by Lemur is only 4 GB and contains information from over 300,000 RefSeq genomes. Lemur and Magnet are open-source and available at https://github.com/treangenlab/lemur and https://github.com/treangenlab/magnet.

## 1 Introduction

The democratization of long-read sequencing has arrived (Marx 2023) and it is now common practice in metagenomic studies due to a combination of higher accuracy, increased affordability, and greater genomic resolution provided by longer reads (Agustinho et al. 2024). One of the most common tasks in metagenomics is to perform taxonomic profiling of a microbial community specific to a host microbiome or environmental microbiome. Existing taxonomic read classification tools such as Kraken 2 (Wood et al. 2019) have established themselves as a *de facto* standard approach for taxonomic read classification and taxonomic profiling with short-read data (when used in combination with Bracken (Lu et al. 2017)). Several new tools have recently been developed to leverage long-reads for taxonomic profiling. Common approaches taken by the developers consist of methods based on *k-mers* (e.g., Kraken 2 (Wood et al. 2019), Sourmash (Irber et al. 2022)), *read-mapping to a succinct index* (e.g., Centrifuger (Song and Langmead 2024), MetaMaps (Dilthey et al. 2019)), *proteins* (e.g., MEGAN-LR (Huson et al. 2018)) and *marker genes* (M Wu and Eisen 2008) (e.g., Melon (Chen et al. 2024), PhyloSift (Darling et al. 2014), MetaPhyler (B Liu et al. 2011)).

Prior studies have highlighted the challenges in benchmarking metagenomic profilers (Sun et al. 2021) and have evaluated the accuracy of new and existing methods for long-read microbiome profiling (Portik et al. 2022); methods explicitly designed for long reads tend to perform better. However, in these prior studies, the experimental evaluation focused primarily on precision and recall. Given that the long-read technologies offer potential for portable and streaming sequence analysis (Quick et al. 2016), it is also important to evaluate scalability and fitness for execution in low-resource environments such as laptops and tablet computers. In particular, depending on the database size and computational requirements, several currently available tools need dedicated server nodes with high RAM (*>* 32 GB) and/or high parallelization capacity to achieve time-to-answer below a few hours (Simon et al. 2019).

To address these issues, we present a two-tool suite consisting of Lemur, a marker-gene-based long-read taxonomic profiler, and Magnet, a genome-based validation tool for confirming the presence and absence of microbial genomes present in a sample. Combined, Lemur and Magnet can run in limited resource settings, such as individual laptops, and yield comparable or superior performance in terms of precision and recall. Our overall contribution hence is threefold: (1) we propose two novel methods for taxonomic profiling that achieve superior precision and portability, (2) we extend the marker gene database to include fungal genes for fungi classification, and (3) we facilitate taxonomic profiling in lightweight compute environments.

## 2 Results

### 2.1 Method overview

An overview of Lemur and Magnet is presented in Figure 1. Both methods require a FASTQ file containing sequencing reads as input. Lemur additionally requires a marker gene (MG) database, whereas Magnet requires a (ideally small) set of genomes. In both cases, the first step is to map the reads to the database using minimap2 (H Li 2018). For our study, we built a new marker gene database with 43 markers for bacteria+archaea and 48 markers for fungi. These markers were validated by prior studies (Nguyen et al. 2014; Shah et al. 2021; Chen et al. 2024; D Kim et al. 2023). For all marker genes, we downloaded the HMM profiles from both the TIPP2 reference package (Shah et al. 2021) and the KofamKOALA database (Aramaki et al. 2020). All HMM profiles used in the studies are provided in the source data. We then built the database using recent versions of NCBI RefSeq: version 221 for both bacteria (329,194 assemblies) and archaea (1,911) and version 222 for fungi (564). A total of 331,669 genome assemblies, including all of their annotated protein sequences and corresponding CDS sequences, were downloaded from NCBI RefSeq. All of the pseudogenes were excluded. We used fetchMGs (v1.2) to extract the selected marker gene sequences with command fetchMGs.pl -m extraction (Sunagawa et al. 2013) and created a mapping between the sequences and their corresponding species rank taxonomic ids with our custom Python script. We used the Emu database creation tool (Curry et al. 2022) for the final step in the database construction for individual marker genes with the command `emu build-database --ncbi-taxonomy`. Finally, individual marker gene databases were concatenated, and a single joint taxonomy mapping was generated for the combined database. The final database was 4.1 GB, containing 3,335,783 sequences.

**Figure 1:**
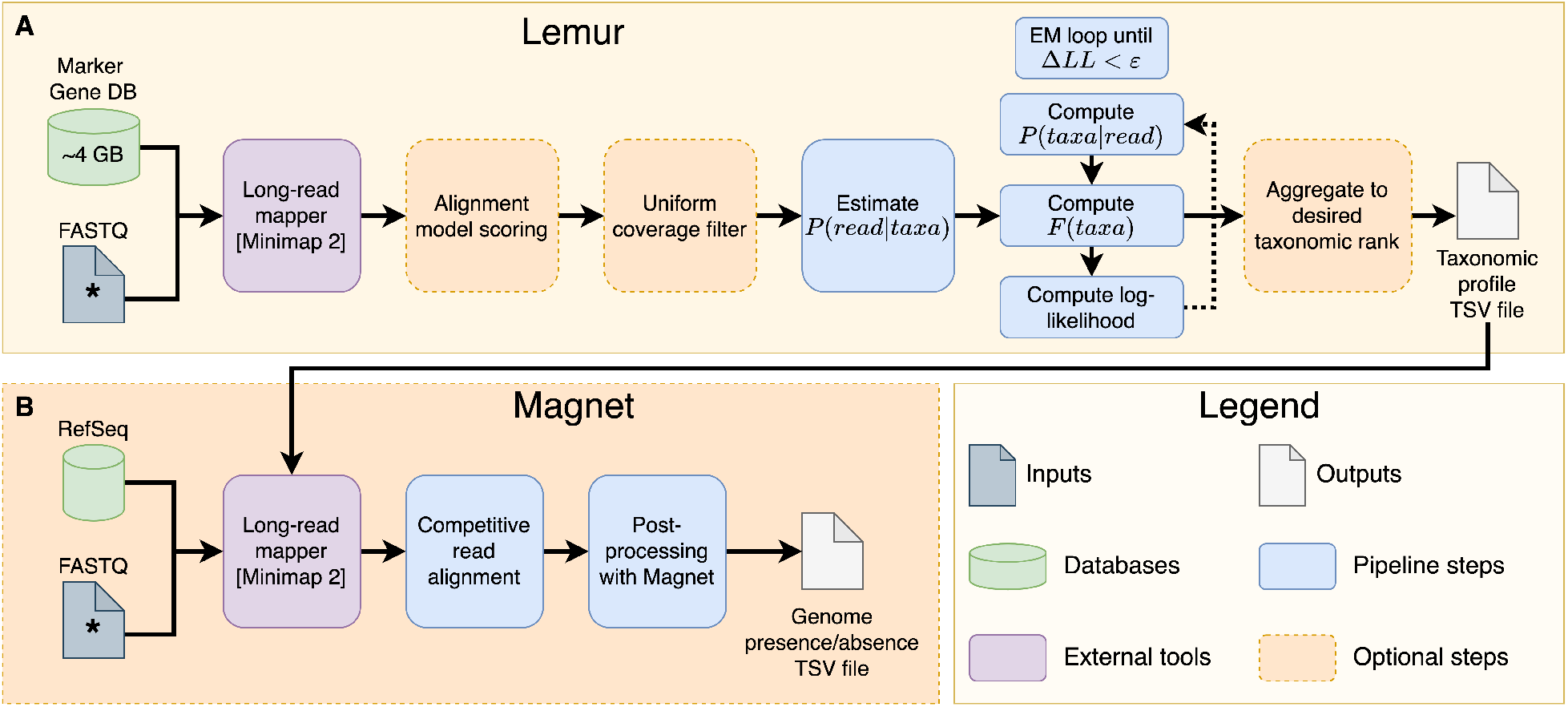
Overview of Lemur **(A)** and Magnet **(B)** pipelines. Input FASTQ files with the * symbol indicate same file. The taxonomic profile table provided to Magnet can come from any taxonomic profiler or classifier, as long as it respects formatting and uses the same set of taxonomic ids.

We conducted a series of evaluations of increased complexity on synthetic, simulated and real data. We compared the performance of our proposed tool Lemur to Centrifuger, Kraken 2, Melon, MetaMaps, and Sourmash. For Lemur results, we also show post-processing with Magnet for improved specificity (Lemur + Magnet).

### 2.2 Prior simulated data

We first evaluated the methods on simulated data from a prior study (Dilthey et al. 2019). On the simulated community of 96 bacterial strains, MetaMaps shows perfect recall at the species level, followed by Kraken 2 with a recall of 0.976. Table 1 contains the full precision and recall data for these five methods on this data. Kraken 2 has high recall but low precision. Melon, Lemur and MetaMaps, all EM-algorithms, each have high recall and moderate to high precision.

**Table 1:** Performance on simulated data from (Dilthey et al. 2019). Recall, precision, and F1 score are defined in methods. Tools listed below the horizontal dashed line focus on the taxonomic classification of reads.

| Method | Recall | Precision | F1 score |
| --- | --- | --- | --- |
| Lemur | 0.951 | 0.703 | 0.808 |
| Lemur + Magnet | 0.927 | **0.950** | 0.938 |
| Melon | 0.963 | 0.929 | **0.946** |
| MetaMaps | **1.000** | 0.862 | 0.926 |
| Sourmash | 0.927 | 0.938 | 0.932 |
| Centrifuger | 0.774 | 0.050 | 0.093 |
| Kraken 2 | 0.976 | 0.055 | 0.104 |

For this analysis we used the MetaMaps results provided in the original manuscript (Dilthey et al. 2019).

### 2.3 ZymoBIOMICS Microbial Standards

We evaluated our tools on the Zymo EVEN and LOG datasets. We evaluated tools on 5 replicate subsamples of the EVEN community (Table 2) at 1% and 75% sampling rates, and 5 replicate subsamples of LOG community (Table 3) at 10% and 75% sampling rates. This was done in order to simulate low and high flow cell usage experiments, respectively.

**Table 2:** Mean performance and standard deviation across 5 replicate runs of all methods on Zymo EVEN, bold values show best performance. Magnet does not report relative abundance, so the L1 error cannot be computed. Tools listed below the horizontal dashed lines (for EVEN 1% and EVEN 75%) focus on the taxonomic classification of reads.

| Dataset | Tool | Recall | Precision | F1 | L1 error (normalized) |
| --- | --- | --- | --- | --- | --- |
| EVEN 1% | Lemur | 0.980 | 0.596 | 0.737 | **0.205** |
|  | Lemur + Magnet | 0.960 | **0.980** | **0.969** | N/A |
|  | Melon | 0.800 | 0.889 | 0.842 | 0.226 |
|  | MetaMaps | **1.000** | 0.909 | 0.952 | 0.340 |
|  | Sourmash | 0.800 | 0.727 | 0.762 | 0.901 |
|  | Centrifuger | 0.800 | 0.571 | 0.667 | 0.413 |
|  | Kraken 2 | **1.000** | 0.589 | 0.741 | 0.321 |
| EVEN 75% | Lemur | **1.000** | 0.669 | 0.801 | **0.144** |
|  | Lemur + Magnet | 0.860 | 0.844 | 0.851 | N/A |
|  | Melon | 0.800 | 0.800 | 0.800 | 0.194 |
|  | MetaMaps | **1.000** | **0.909** | **0.952** | 0.329 |
|  | Sourmash | 0.800 | 0.727 | 0.762 | 0.893 |
|  | Centrifuger | 0.800 | 0.571 | 0.667 | 0.415 |
|  | Kraken 2 | **1.000** | 0.556 | 0.714 | 0.321 |

**Table 3:** Mean performance and standard deviation across 5 replicate runs of all methods on Zymo LOG, bold values show best performance. Magnet does not report relative abundance, so the Spearman’s *ρ* cannot be computed. Tools listed below the horizontal dashed lines (for LOG 10% and LOG 75%) focus on the taxonomic classification of reads.

| Dataset | Tool | Recall | Precision | F1 | Spearman’s $\rho$ |
| --- | --- | --- | --- | --- | --- |
| LOG 10% | Lemur | 0.500 | 0.301 | 0.376 | **0.984** |
|  | Lemur + Magnet | 0.360 | **0.683** | 0.468 | N/A |
|  | Melon | 0.420 | 0.587 | **0.488** | 0.960 |
|  | MetaMaps | **0.960** | 0.009 | 0.017 | 0.976 |
|  | Sourmash | 0.500 | 0.615 | 0.550 | 0.940 |
|  | Centrifuger | **0.960** | 0.002 | 0.005 | 0.906 |
|  | Kraken 2 | 0.760 | 0.249 | 0.375 | 0.910 |
| LOG 75% | Lemur | 0.700 | 0.219 | 0.333 | 0.971 |
|  | Lemur + Magnet | 0.580 | **0.852** | **0.690** | N/A |
|  | Melon | 0.600 | 0.592 | 0.595 | 0.974 |
|  | MetaMaps | **1.000** | 0.002 | 0.005 | **0.985** |
|  | Sourmash | 0.500 | 0.352 | 0.413 | 0.920 |
|  | Centrifuger | 0.960 | 0.001 | 0.003 | 0.909 |
|  | Kraken 2 | 0.720 | 0.246 | 0.367 | 0.903 |

**Table 4:** 9 Genera predicted by Kraken2/Melon.

| Genus | Isolation Source | Citation |
| --- | --- | --- |
| *Anaeropeptidivorans* | Biogas fermenter | Köller et al. 2022 |
| *Caloranaerobacter* | Marine thermal vents | Wery et al. 2001 |
| *Caminicella* | Marine thermal vents | Alain et al. 2002 |
| *Vallitalea* | Marine sediment | Lakhal et al. 2013 |
| *Caproiciproducens* | WW treatment plant | BC Kim et al. 2015 |
| *Lachnoanaerobaculum* | Human/Gut feces | Hedberg et al. 2012 |
| *Monoglobus* | Human/Gut feces | CC Kim et al. 2017 |
| *Solibaculum* | Human/Gut feces | Sakamoto et al. 2021 |
| *Vescimonas* | Human/Gut feces | Kitahara et al. 2021 |

On the EVEN community, Lemur showed high recall matched by MetaMaps and Kraken 2. We note that Lemur can achieve full recall without restricting evaluation to bacteria-only, suggesting that it is capable of accurately evaluating both kingdoms jointly. Additionally, the benefit of polishing the abundance profiles with Magnet is supported by an increase in precision particularly in low-coverage scenarios where false-positives are more likely. Finally, in both low and high-coverage settings, Lemur showed lowest L1 error.

On the LOG community, recall of all tools except MetaMaps drops as some species are present at such a low level that their expected genome copy number is less than 0.05. In particular, marker-gene-based tools Melon and Lemur have recall lower than MetaMaps, Centrifuger and Kraken 2 which can utilize the information from across the whole genome (Table 3). By contrast, the combination of many low-abundance taxa and low-coverage creates a large number of false positives as shown by the lower precision of most tools (Table 3). However, Magnet does well with identifying false positives and improves the precision by nearly 0.6 over Lemur alone. This indicates that the tools retain the ability to make confident species level calls in extremely low-abundance settings (Table 3).

Additionally, Lemur shows strong performance on Spearman’s *ρ* (see Section 4.7 for details) indicating that its abundance estimates are broadly accurate vis-a-vis ordering of the microbes within a sample. Further, the reduction in variance of *ρ* in the higher-coverage sample is reassuring since this is an expected outcome of the additional data (Table 3, rightmost column).

### 2.4 Simulated metagenome

The simulated metagenome dataset is intended to present the tools with a more challenging setting, with more diverse taxa present, but where ground truth is *known*. At the species level, Sourmash and Melon show the highest recall, closely followed by Lemur (Figure 2A). Post-processing with Magnet reduces recall but improves precision, in particular achieving the highest precision among all tools at both species and genus levels (Figure 2A-B). We also note that Sourmash and Lemur have high precision at the genus level (Figure 2B), indicating that most false positive species-level calls made by these tools come from the correct genus.

**Figure 2:**
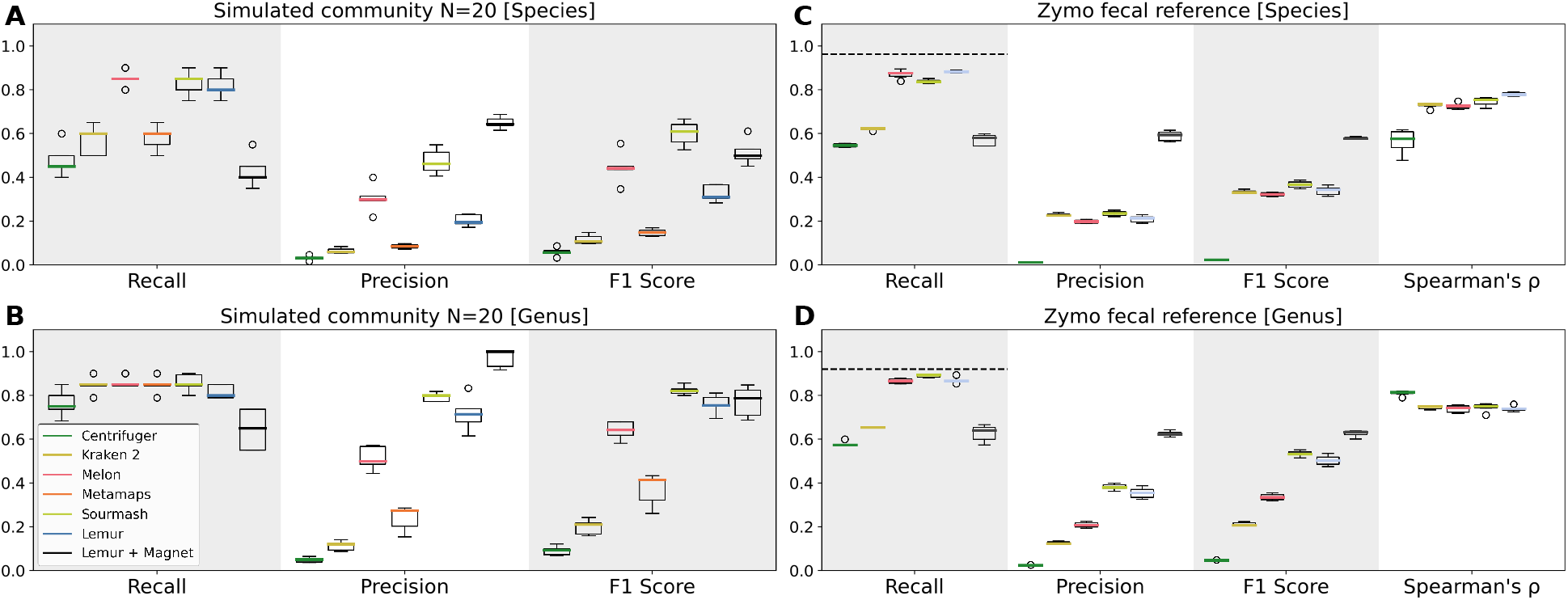
Performance of all methods on a simulated metagenomic community (A, B) and Zymo fecal reference (C, D). Panels show recall, precision, F1 score and Spearman’s *ρ* (**C, D**) for each of the tools. Dotted lines (C, D) indicate maximum recall based on the NCBI RefSeq v222 composition.

### 2.5 ZymoBIOMICS Fecal Reference

Next, we evaluated the performance of all tools on the ZymoBIOMICS fecal reference samples (Figure 2C-D). We did not include MetaMaps in this evaluation due to computational constraints. Lemur and Melon are the most sensitive tools at the species level, closely followed by Sourmash (Figure 2C), at the genus level Sourmash has the highest recall with Lemur and Melon matching closely (Figure 2D). However, the precision of Sourmash and Lemur is higher than Melon at both ranks. As with the previous datasets, post-processing with Magnet improves precision, although it imposes a penalty on recall. The tool combination maintains the highest F1 score (Figure 2C-D). Lemur also has the highest *ρ* at the species level. At the genus level, Centrifuger has the highest Spearman’s *ρ*, although as noted in section 4.7, the calculation in this case ignores false positives and false negatives (Figure 2C-D, rightmost panels).

### 2.6 Human gut metagenome

Finally, species and genus level calls made by Lemur were evaluated on stool sample from a healthy donor (CY Kim et al. 2022). We note that the most abundant phyla identified by Lemur (Figure 3) match those in the prior literature. In particular *Actinomycetota, Bactriodota, Bacillota*, and *Pseudomonadota* are well-represented phyla (Figure 3P). Additionally, *Faecalibacterium prausnitzii, Bacteroides ovatus, Parabacteroides distasonis*, and *Alistipes onderdonkii* (Figure 3S) are all supported by the co-assemblies from the original study. Other bacterial species identified by Lemur such as *Roseburia faecis* and *Phocaeicola vulgatus*, have all been previously isolated from human feces and can be considered putative true-positives (Almeida et al. 2021; Mancabelli et al. 2024; J Li et al. 2014; Costea et al. 2017)(Figure 3S).

**Figure 3:**
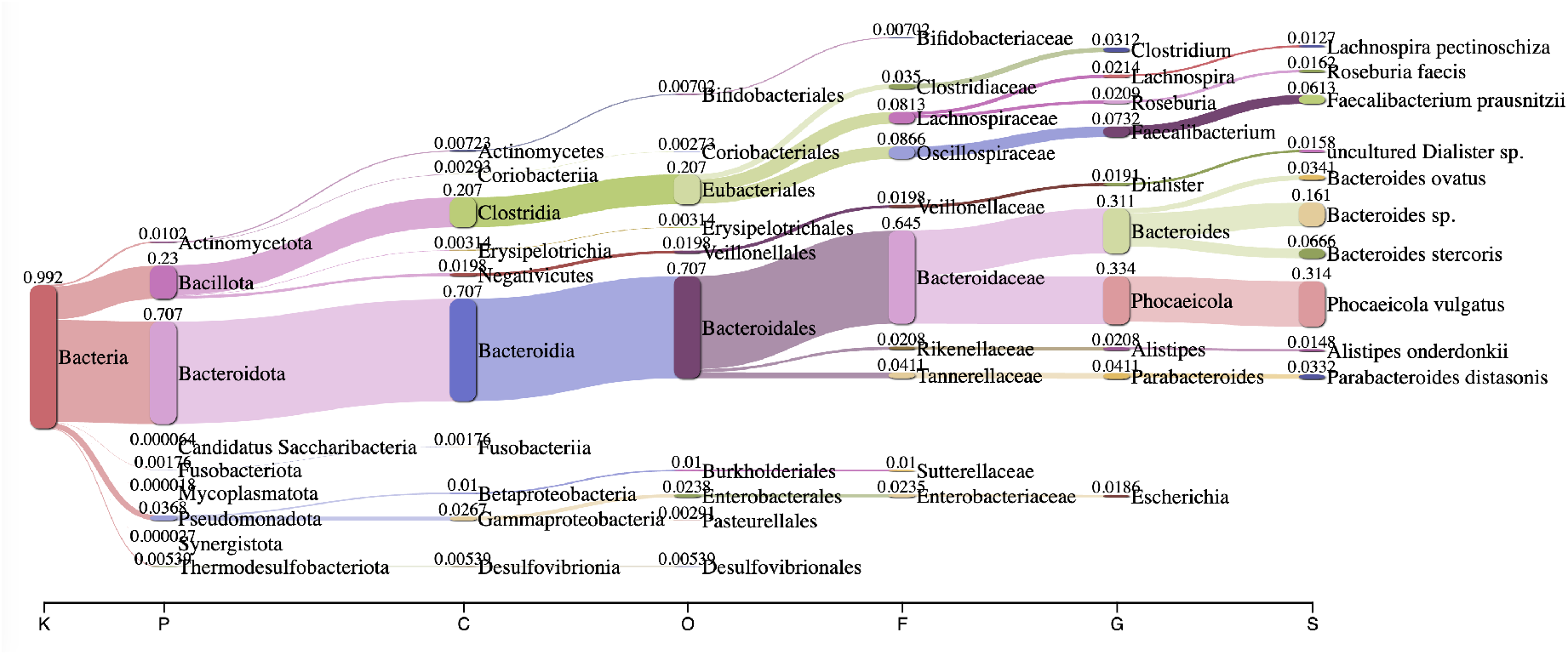
Sankey plot of major taxonomic groups inferred by Lemur from a stool sample from a healthy human donor.

To perform a deeper dive, we also investigated the concordance between the species and genus level calls across the tools. One genus level call that is uniquely inferred by Lemur corresponds to *Massilimicrobiota*. A recent isolate from the human gut belonging to the *Massilimicrobiota* genus has been identified (Tall et al. 2019), indicating that this call is a plausible true positive. Conversely, 9 genera were identified by Kraken 2 and Melon but *not* Lemur. Those include several genera whose reported isolation sources are distant from the human gut, including a biogas fermenter, two from marine thermal vents, marine sediment, and a wastewater treatment plant. The remaining four genera were previously found in the human gut. The list of genera, their isolation source, and relevant citations is contained in Section A.2.1 of the Supplement.

While this is not a conclusive experiment to assess false positives, it represents a sensible evaluation of a new method and a step towards improved precision in challenging settings.

### 2.7 Chicken gut metagenome

Additionally, we have analyzed a chicken gut metagenome spanning four major sections of the intestinal tract: duodenum, jejunum, ileum, and colorectum (last two shown in Figure 4A, B). Similar to prior studies, our results highlight an increase in microbial diversity along the intestinal tract (Y Zhang et al. 2022; P Huang et al. 2018). Furthermore, we noted that the samples from foregut (duodenum, jejunum, ileum) were dominated by a few abundant taxa from the *Lactobacillacea* family (Figure 4A). At the same time, the hindgut showed a more even abundance distribution across a broader set of taxa. These observations are consistent with results obtained from metagenomic assembly in the original study (Y Zhang et al. 2022).

**Figure 4:**
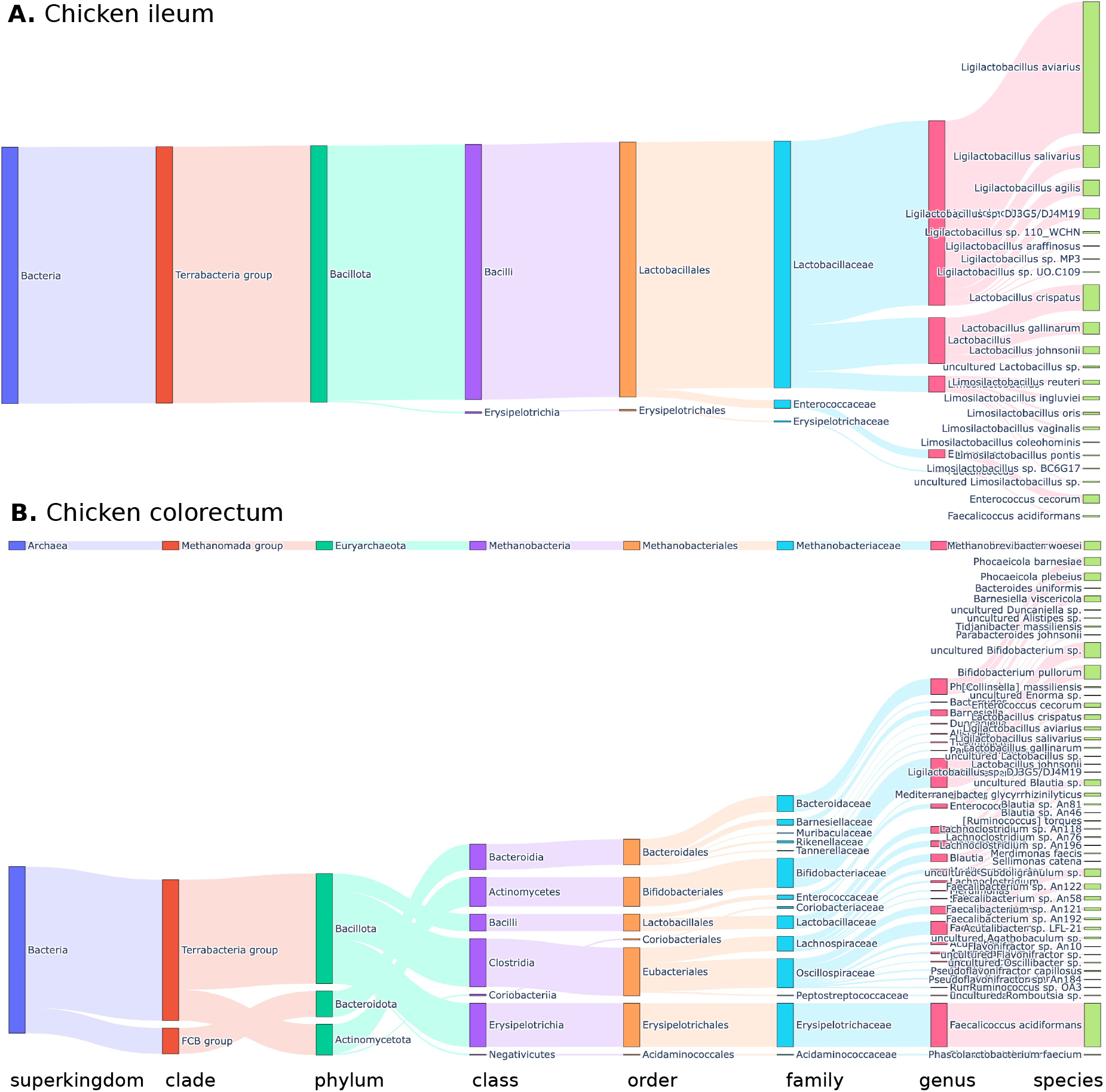
Sankey plot of major taxonomic groups inferred by Lemur from a chicken gut metagenome of two intestinal compartments. Top panel corresponds to the ileum (**A**) and bottom panel represents the colorectum (**B**).

The colorectum shows the most diverse set of taxa (Figure 4B), including a single archaeal species in class *Methanobacteriota*. Notably, among the 337 assemblies from the original study (Y Zhang et al. 2022), there is exactly one archaeal genome. Analogously to the human gut metagenome example, this data does not definitively assess the accuracy of Lemur; however, it suggests that the inferred taxonomic profiles match closely with the prior literature (P Huang et al. 2018; Y Zhang et al. 2022).

### 2.8 Computational performance

The computational performance of Centrifuger, Kraken 2, Melon, MetaMaps, Sourmash, and Lemur for each of the four main datasets is captured in Figure 5. Sourmash, Melon and Lemur have their total RAM requirement consistently under 32 GB, with Lemur requiring more RAM than Melon and Sourmash (Figure 5G-J). The RAM requirement for MetaMaps exceeds 300 GB, as indicated by an asterisk (*) on the plot. CPU time requirements for the tools are consistent on small and medium-sized datasets (Figure 5A-E), with Kraken 2 requiring the least time, followed by Lemur. Notably, Sourmash and MetaMaps require at least an order of magnitude more CPU time (Figure 5A-E, log-scale y-axis). Finally, post-processing with Magnet doesn’t incur much additional cost for small and medium datasets (Figure 5A-E, time shown for serial execution of Lemur and Magnet). On the large and diverse data (Figure 5F) the overall trend remains the same with Kraken 2 being the fastest tool, followed by Lemur. However, in this case post-processing with Magnet takes a significant portion of total time due to richer microbial composition of the dataset.

**Figure 5:**
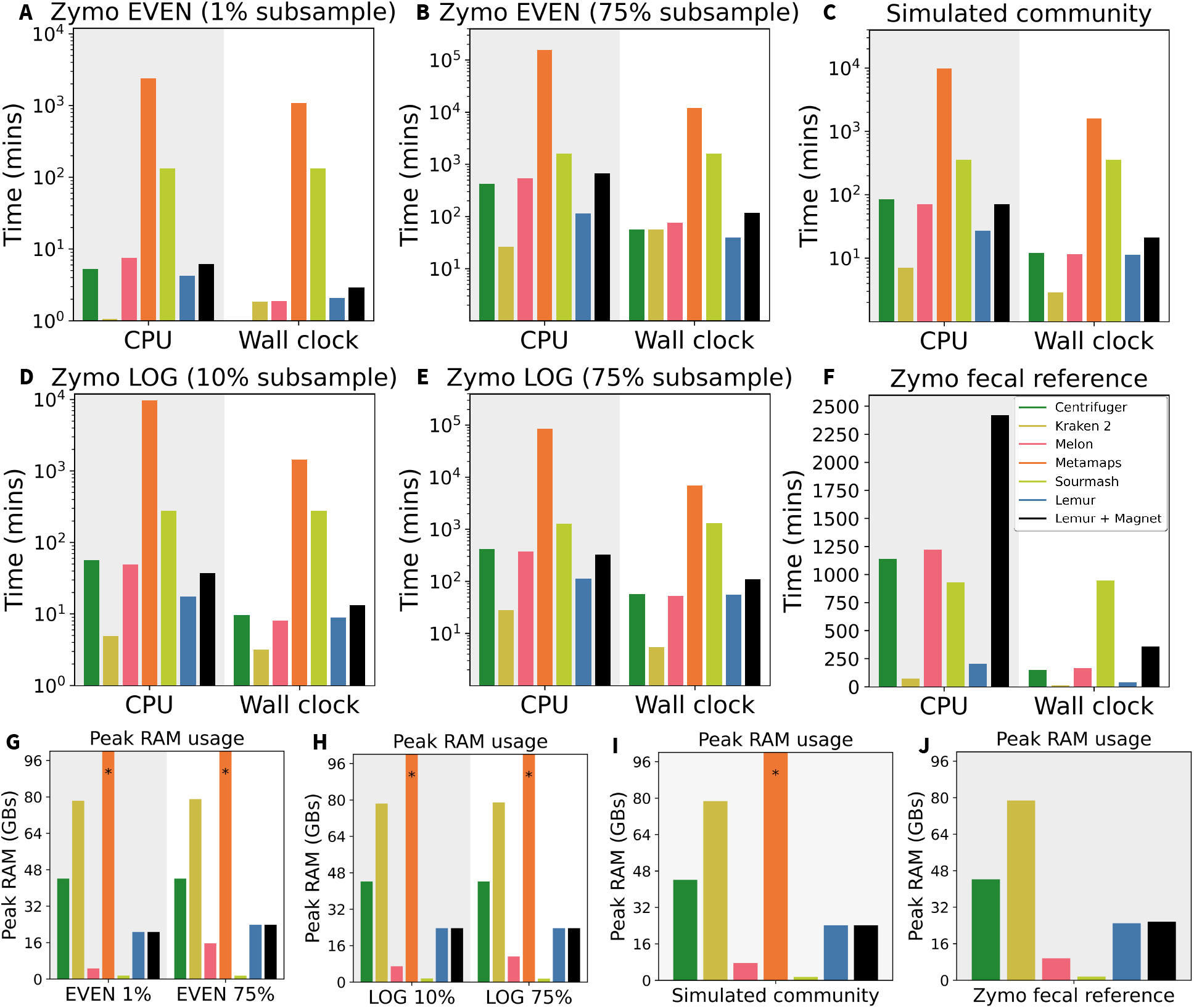
Computational performance metrics evaluated on Zymo EVEN (**A, B, G**), Zymo LOG (**D, E, H**), simulated metagenome (**C, I**), and Zymo fecal reference (**F, J**) datasets. Top six panels show CPU and wall clock times (**A**-**F**), and bottom four panels show peak RAM usage (**G**-**J**).

Overall, Kraken 2 is well-known for its ultrafast processing of metagenomic samples but at the cost of high RAM requirements for larger databases. Lemur has the second lowest CPU time requirement and performs comparably to Melon in terms of wall clock time for small and medium-sized datasets. For large datasets, Lemur appears to have a larger CPU time advantage over the other methods. Magnet incurs a moderate additional cost for post-processing of the results and hence is an attractive option for improved precision of the analyses.

Lastly, the feasibility of using Lemur for taxonomic profiling directly on a user laptop was evaluated using two datasets of different sizes. Both were run with Lemur on a MacBook Pro (macOS Sonoma 14.2.1) having a 2.3 GHz Intel Core i9 and 32 GB of RAM using 4 threads. The first dataset was a 3 GB Zymo LOG sample which ran in 13.56 mins (CPU time: 21.99 mins) with peak RAM usage of 17.09 GB. The second was a 77 GB Zymo Fecal Reference sample, which ran in 73.45 mins (CPU time: 206.66 mins) with peak RAM usage of 23.05 GB. Given the real-time and portable nature of long-read sequencing devices, this demonstrates an essential element of compatibility with many envisioned use cases in the future.

### 2.9 F1 score per CPU time unit

For the combination of the datasets and tools evaluated, we have also investigated the median achieved F1 score as a function of the CPU time required to analyze the data. We note that for Zymo EVEN 1%, Zymo LOG 75%, and Zymo fecal reference data Lemur + Magnet achieve the highest total F1 score while requiring an amount of CPU time comparable to other tools (Figure 6A, E, F). For the cases where Lemur + Magnet do not achieve the highest F1 score (Figure 6B-D) the tools that do require at least an order of magnitude more CPU time.

**Figure 6:**
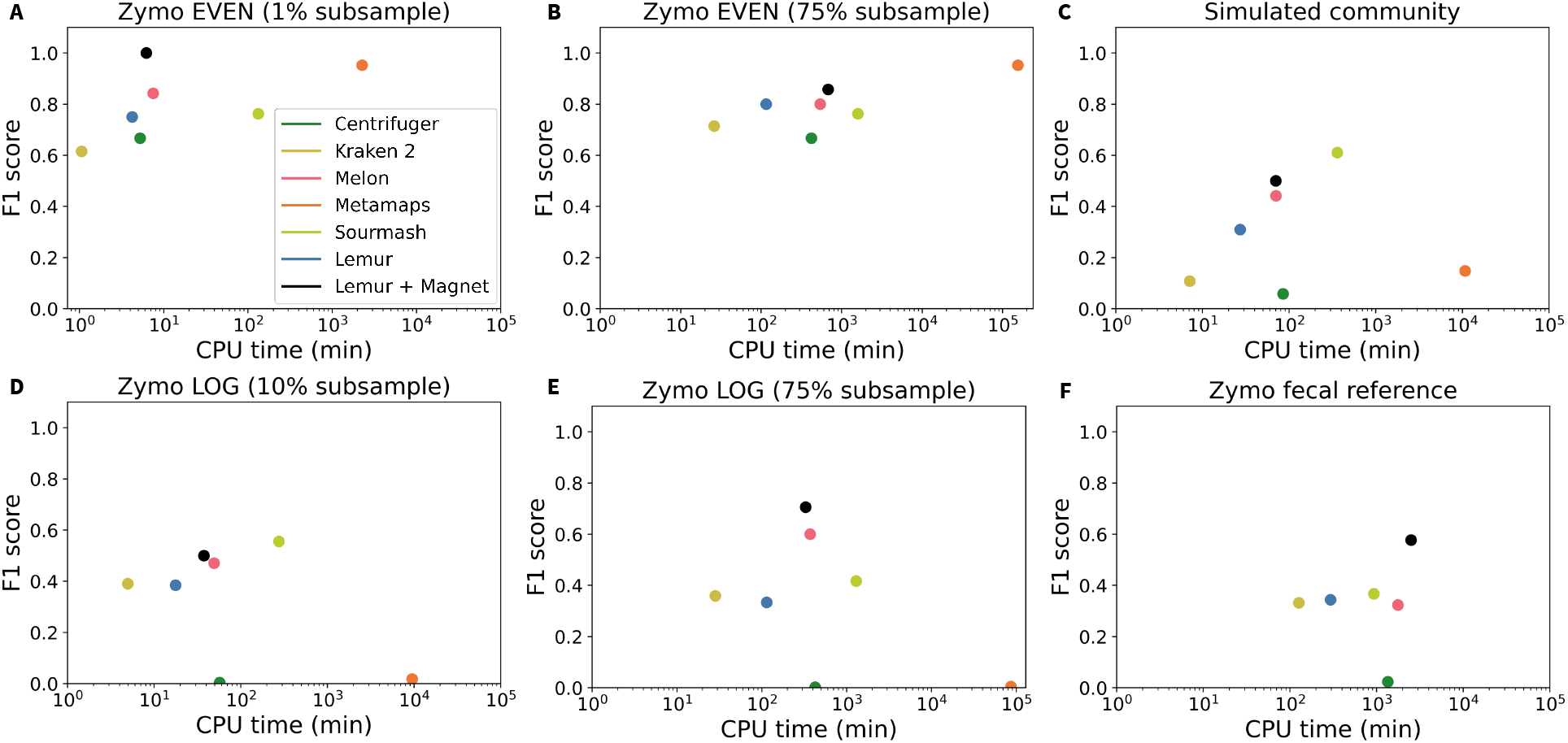
Median F1 score achieved by the tools as the function of required CPU time for processing (on logarithmic scale) for the datasets used in the study.

## 3 Discussion

Lemur and Magnet represent a novel computational tandem tailored for long-read taxonomic profiling of metagenomes. Lemur exhibits competitive performance by most standard metrics; when paired with Magnet, particularly in the presence of low-abundance or low-coverage data, it can improve precision by detecting and filtering out many false positive calls. Our results indicate that Lemur can efficiently process large datasets within minutes to hours in limited computational resource settings. Furthermore, consistent RAM usage below 32 GB makes it an attractive software for laptops and lightweight workstations. This reduced resource need does not come at the cost of accuracy. Lemur and Magnet combined offer high recall and precision on the experimental datasets included in our evaluation. Our comparative analysis of human gut microbiomes indicates that Lemur produces a taxonomic profile consistent with the gut microbiome while avoiding outlier species calls.

Inspired by previous marker-gene-based approaches (B Liu et al. 2011), relying on a wide pool of single-copy universal marker genes allows Lemur to achieve high recall and relative abundance estimation accuracy while using only a small portion of the input data. These markers cover all bacteria but only a fraction of any given genome. In contrast, Magnet starts with the set of genomes identified by Lemur and evaluates the read-alignment quality and coverage distributions across the genome to make a maximally informed call about whether the putative genome is actually in the sample. Additionally, the inclusion of fungi into the database broadens the scope of the tool and allows for a more comprehensive characterization of metagenomes.

As with any computational method, Lemur and Magnet have limitations that vary by use case. Reliance on bacterial marker genes necessarily implies it cannot generalize to viral genome classification. Also, while Lemur and Magnet can filter out false positives in low-abundance/low-coverage settings, the reliance on the marker genes makes it less sensitive than alternatives like Kraken 2 or MetaMaps, which use all long reads and complete genomes. Third, the nature of the EM algorithm employed means that it is by necessity a closed-reference method, and thus, a bacteria from a novel, *i*.*e. out-of-database* family will necessarily be missed by Lemur. Finally, marker gene methods inherently lack the resolution to perform strain-level classification or other sub-species analysis. However, the pairing with Magnet suggests one approach for sub-species inference that could change this in the future.

Overall, our results on simulated, synthetic, and real datasets provide experimental support that the combination of Lemur and Magnet represents an efficient and accurate taxonomic profiling workflow explicitly designed for long-read sequenced metagenomes. In future work, we intend to expand our benchmark to include additional tools (J Kim and Steinegger 2024; Shaw and Yu 2023; Peres da Silva et al. 2024) and datasets to provide a more comprehensive landscape of long-read taxonomic profilers and classifiers, including analysis of how the performance of these tools varies with differences in reference database compositon (Nasko et al. 2018). Furthermore, we acknowledge that while our study focused on taxonomic profiling and binary presence and absence metrics for taxa, several of the considered methods (Kraken 2, Centrifuger, MetaMaps) are in fact *metagenomic read classifiers*. Thus, a separate benchmark that focuses on the percentage of classified reads and the proportion of correctly classified reads is warranted to properly assess the accuracy of these methods for the specific computational tasks they were designed for.

## 4 Methods

### 4.1 Lemur

Lemur estimates a relative abundance taxonomic profile using an Expectation-Maximization (EM) framework, similar to the 16S profiler Emu (Curry et al. 2022). The EM algorithm proceeds in two steps. First, in the E-step, we compute the probability *P* (*t*|*r*) of taxon *t* being in the sample given an observed read *r*, which uses an abundance prior *F* (*t*) and Bayes rule: 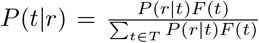, where *P* (*r*|*t*) is the probability of observing read *r* originating from taxon *t*. Second, in the M-step, the abundance vector *F* (*t*) is re-estimated using *P* (*t*|*r*) for all reads *r* in the dataset as 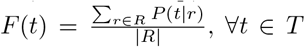. Finally, we compute the total log-likelihood

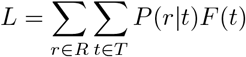

and the change in log-likelihood Δ*LL* from the prior step *L*^*′*^ as Δ*LL* = *L − L*^*′*^. These steps are repeated until Δ*LL* < *ε*, where *ε* is a user-defined threshold with a default value of 0.01 (see blue cycle of Figure 1A). After convergence, Lemur removes any taxonomic assignments below a user-specified threshold (default: 1 / number of reads that map to marker genes) and then runs a final iteration of the EM algorithm to re-estimate *F* (*t*).

The EM algorithm begins by initializing *F* (*t*) to the uniform distribution and initializing *P* (*r*|*t*) for each read and taxon pair (*r, t*). Lemur implements several options for calculating *P* (*r*|*t*). The fastest and default option is to take the alignment score output by minimap2 and normalize it: *P* (*r*|*s*) = *AS*(*r, s*)*/*2*L* where *AS* is the alignment score from the minimap2 SAM file and *L* is the alignment length. Lemur aggregates the scores for all sequences assigned the same taxonomic label, retaining the score from the most likely alignment, setting *P* (*r*|*t*) = max_*s∈t*_ *P* (*r*|*s*) (as there can be multiple genomes corresponding to the same taxon). Although not explored in this study, Lemur implements two other approaches for estimating *P* (*r s*), including the CIGAR-based approach from Emu (Curry et al. 2022) and a Markov chain model of sequence alignment.

Additionally, Lemur can employ a uniform coverage (UC) filter before the EM-steps to eliminate taxa from the taxonomic profile if the read-coverage pattern across the marker genes significantly deviates from the expected value. The goal is to improve Lemur’s precision when sequencing depth is high, and coverage is uniform (so taxa with non-uniform coverage may be false positives). Specifically, let *G* be the number of marker genes. Let *N* be the number of reads that map to a particular taxon. Let *X* denote the random variable corresponding to the number of distinct genes with at least one read aligned to them. Let *X*_*i*_ be the indicator variable for the event that a gene *i* is covered by a read. Then, the expected number of genes hit is

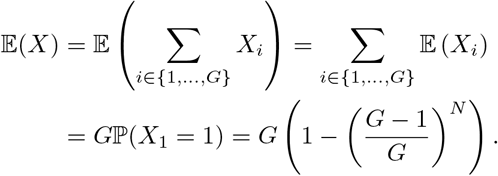

The first equality is by linearity of expectation, and since the event of a gene being hit by at least one read is complementary to all reads aligning to one of the other *G* – 1 genes, it follows that the desired equality holds (we treat individual read to gene alignments as independent events). Furthermore, we note that:

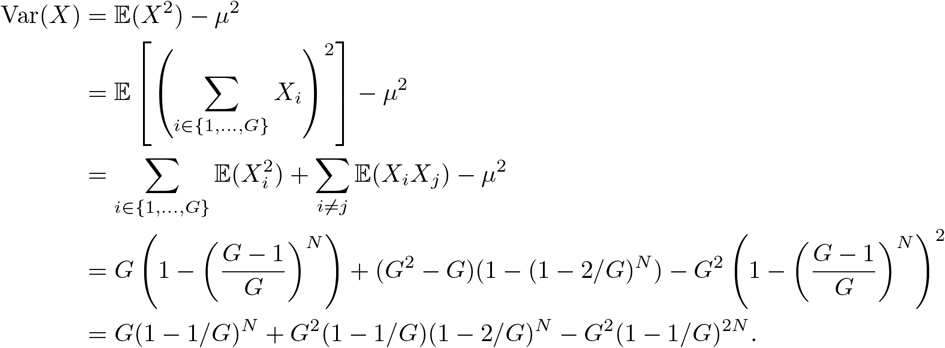

Again, the first expansion is due to the linearity of expectation. Then we note that the random variable 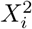 has the same expected value as *X*_*i*_, while ℙ (*X*_*i*_*X*_*j*_ = 1) is given by 1 *–* ℙ (*V*_*k≠i,j*_ *X*_*k*_ *= 1)* which is simply 1 – ((*G* –)/*G*)^*N*^

Finally, the actual number of genes hit, denoted *G*_*hit*_, is also computed directly from the data. If *μ − G*_*hit*_ ≤ 3*σ*^2^ and *G*_*hit*_*/μ* ≤ 0.7, the *taxon* is removed (filtered) from further consideration.

As a post-processing step, the taxonomic profile, along with the original reads, can be provided to Magnet as input for genome presence and absence evaluation. Magnet employs a competitive alignment strategy designed to differentiate closely related genomes.

### 4.2 Competitive read alignment with Magnet

The goal of Magnet is to detect and remove potential false positives by performing competitive read alignment leveraging all of the reads mapped against the entire reference genome (rather than just the marker gene reads and marker genes used by Lemur). As input, Magnet requires reads as well as a taxonomic abundance profile (estimated from the input reads e.g. using Lemur). For each candidate species identified in the profile, Magnet downloads the highest ranking genome from NCBI RefSeq and Genbank databases, where the ranking is determined by (1) representative genomes in RefSeq, (2) complete genomes in RefSeq, (3) other genomes sorted by assembly level (complete, chromosome, scaffold, contig).

Once the genomes are downloaded, Magnet constructs a dissimilarity matrix by calculating pairwise average nucleotide identity (ANI) between all candidate sequences using FastANI (Jain et al. 2018). Magnet then performs agglomerative clustering with scikit-learn (Pedregosa et al. 2011) using a threshold of 0.05. For each cluster, one representative genome is selected. The representative genome is picked to maximize the sum of pairwise ANI to all other genomes in the cluster. If a cluster contains complete genomes, the representative genome is required to be complete, otherwise it can be at any assembly level.

The reads from the input dataset are aligned to all cluster representative genomes at once using minimap2. Magnet calculates the observed breadth and depth of coverage for the alignment with `samtools coverage` for two cases: (1) including all alignments with MAPQ ≥ 1 and (2) including only primary alignments with MAPQ ≥ 20. For both cases, Magnet calculates the expected breadth of coverage based on the abundance and the coverage score, defined as the ratio of the observed breadth of coverage and expected breadth of coverage (Balaji et al. 2023). The coverage score is used to measure the uniformity of the alignment distribution along the reference genome. Magnet generates the consensus genome for each cluster based on alignments and then estimates ANI between it and the reference genome.

Lastly, Magnet marks species as present or absent. If the consensus versus reference ANI is greater than 0.97 and the breadth of coverage for high MAPQ primary alignments is greater than 0.2, the species is determined to be present. The species is also marked as present if ANI is greater than 0.7 and the *reduction* in coverage score and breadth of coverage between all alignments and high MAPQ primary alignments is less than or equal to 0.1. Otherwise, the species is marked as absent. The Magnet presence and absence calling process is based on both the ANI between consensus and the aligned region reference, and coverage and uniformity decrease after excluding low MAPQ alignment. For species that are truly present in the sample, excluding low MAPQ alignments has a limited effect on the breadth of coverage or coverage score. On the other hand, excluding low MAPQ alignments usually causes a significant breadth of coverage or coverage score reduction for likely false positives, as alignments to the false positive species usually have a lower average MAPQ. The implemented thresholds were chosen based on empirical evidence and simulation-based testing.

### 4.3 Method Comparison

We compared the performance of Lemur (v1.0.1) to taxonomic classification tools Centrifuger (v1.0.0) (Song and Langmead 2024), Kraken 2 (v2.1.3) (Wood et al. 2019), Melon (v0.1.0) (Chen et al. 2024), MetaMaps (commit: 633d2e0; Oct 10, 2023) (Dilthey et al. 2019), and Sourmash (v4.8.2) (Irber et al. 2022). Lemur was evaluated both on its own and in combination with Magnet (Lemur + Magnet). Melon doesn’t include any fungal references in its database and thus cannot classify fungi. Therefore, for two datasets that include fungi in the ground truth, we report Melon and Lemur results on bacterial species only in addition to results using complete ground truth. Exact commands used for running each tool are provided in Section A.1.1 of Supplement.

### 4.4 Synthetic and simulated datasets

#### 4.4.1 Simulated data from (Dilthey et al. 2019)

Our first dataset is a simulated dataset from (Dilthey et al. 2019), with 96 bacterial strains. After validating the metadata, we confirmed that 94 out of the specified 96 strains have a corresponding species representative in the RefSeq database. The dataset contains 200,114 reads with an average length of 4,997bps and has a total size of 1.9 GB. Additional simulation details are provided in the original manuscript (Dilthey et al. 2019).

#### 4.4.2 Zymo EVEN & Zymo LOG

The Zymo EVEN data sets were constructed from the ZymoBIOMICS Microbial Community Standard (D6300) sample, which consists of a DNA mixture of 8 bacterial species at an even total DNA abundance of 12% and 2 fungi at 2% DNA abundance. The Zymo LOG datasets were constructed from the ZymoBIOMICS Microbial Community Standard (D6310) sample, which consists of a DNA mixture of 8 bacterial species in a 10-fold dilution series by total DNA abundance and 2 fungi. Species used in D6310 are identical to those in D6300, although the abundances in the latter are intentionally given a heavily skewed distribution, with several low-abundance taxa present.

Original sequencing data from a Nanopore GridION device with R9.4 chemistry produced by Loman lab (Nicholls et al. 2019) was downloaded from GitHub. Experiments to understand the effect of coverage level were conducted by sub-sampling the reads with the `seqkit sample` command (Shen et al. 2016) by setting the proportion of sampled reads to 1% (Median # reads: 34.8k, avg. len.: 4006bp, avg. size: 272MB) and 75% (2.61m, 4012bp, 20GB) for Zymo EVEN and to 10% (367k, 4368bp, 3.1GB) and 75% (2.75m, 4372bp, 23GB) for Zymo LOG. For each sampling threshold, this process was repeated five times (setting the random seed parameter from 1 to 5) to create 5 replicate datasets.

### 4.4.3 Simulated metagenome

We simulated a challenging metagenome using 50 species with representative genomes deposited in RefSeq, with metagenome assembled genomes (MAGs) available and having an assembly-quality of at least ‘scaffold’ were selected. 5 replicate datasets where then created by randomly sampling 20 genomes spanning 20 distinct species, and 18-20 distinct genera. Reads were simulated from these genomes using pbsim3 (Ono et al. 2022) with a quality-score based ONT HMM profile and a mean read-length of 4,050bps (S.D: 1,000bp). Reads were then combined into simulated metagenomic samples (avg. size: 6GB). The exact pbsim3 command is provided in Section A.1.2 of the Supplement.

### 4.4.4 Zymo Fecal Reference with TruMatrix

Our final dataset is sequencing data generated on a Pacific Biosciences Sequel IIe device from the Zymo-BIOMICS fecal reference samples. These data along with reference taxonomic abundance profiles were downloaded from ZymoBIOMICS and were based on a separate, curated analysis based on multiple rounds of Illumina shotgun sequencing. A total of 6 samples were analyzed ranging in size from 54 to 105GB (avg. read len.: 6973bps, avg. size: 75.8GB).

This dataset is challenging to evaluate because some community members are listed as unknown at the species and, in several cases, at the genus level. Additionally, several species are listed as ‘uncultured (genus)’ which is problematic because this is also a possible species-label that the tools can output, although in that case it would represent something less than a clear true positive. To avoid this ambiguity, for this dataset all such taxa were excluded from evaluation. The filtered abundance profile containing 180 bacterial species and 1 archaeal species spanning 78 genera was used as the ground truth.

### 4.5 Gut sample from healthy donor (SRR17687125)

For a real metagenomic dataset, a gut metagenomic sample obtained from a Korean donor reportedly in good health and with an omnivorous diet was used (CY Kim et al. 2022). The sample was sequenced on a Pacific Biosciences Sequel II device and contained 2.02M reads with an average read-length of 14,670bp. While no ground truth is available for this sample, the set of the assembled genomes from the original study (CY Kim et al. 2022) was used as a proxy for organisms present in the sample. False-positive calls were not assessed, only the recovery of taxa from assembled genomes as well as the consensus among different taxonomic profilers in the study. This experiment was partly intended as a sanity check for Lemur to see that it produces a plausible gut community without any highly improbable microbes.

### 4.6 Chicken gut metagenome

Additionally, 6 chicken gut metagenome samples spanning different sections of the intestinal tract were analyzed. The samples were sequenced on a Pacific Biosciences Sequel II device and contained 521,403– 4,282,198 reads (avg.: 2,615,037) with an average read-length of 15,989bp. Similarly to the human gut metagenome no ground truth is available for this dataset, but a set of high quality MAGs is available from the original study that produced the data (Y Zhang et al. 2022).

### 4.7 Method evaluation

For all the tools and datasets with known ground truth, we report recall, precision, and F1 score. Precision and recall are defined at the genus and species levels and, in each case, over the set of taxa that are known or estimated to be present above a threshold of 1e-12 (with the exception of Zymo EVEN experiments, where the threshold was set to 1e-3). This was applied to convert abundance profiles for presence and absence.

For tools that provide relative abundance estimates or can be used to derive relative abundance, we also evaluate normalized L1-loss and Spearman’s rank correlation coefficient *ρ*. For datasets with < 20 ground truth species and even abundance distribution we use normalized L1-loss defined as 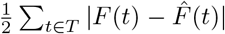, where *F* is actual and 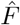 is estimated abundance. L1-loss is less informative for datasets with large numbers of taxa or with an uneven distribution, so we use Spearman’s *ρ* as an alternative metric of correspondence between *F* and 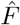. In this case, it is defined over the set of species *t* such that both *F* (*t*) > 0 and 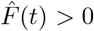 (so methods are not punished for false positives or false negatives in the Spearman’s *ρ*).

All methods except for MetaMaps were ran on a server running Red Hat Enterprise Linux (v8.9) with an AMD EPYC 7742 64-core processor and 128 GB of RAM. All server runs were executed with 8 threads to emulate a laptop-like environment. No additional restrictions were placed on the RAM available to the tools. MetaMaps runs were performed utilizing 20 (for 1% and 10% subsamples) and 30 (for 75% subsamples) threads on servers with 4 Intel Xeon Gold 6150 (18-core) and 4 Intel Xeon Gold 6240 (18-core) processors and 1.5 TB of RAM running CentOS 7.9.2009 Linux. Separate experiments on laptop hardware are described later.

## 5 Data access

Simulated metagenomic sequencing reads and subsampled reads from Zymo EVEN and LOG microbial standards are available in Box. Prebuilt Lemur database is available on Zenodo [DOI:10.5281/zenodo.10802545].

## 7 Acknowledgements

The authors would like to thank Austin Marshall for his feedback on the manuscript and Daniel Portik and Dan Nasko for their dataset recommendations. This work is supported in part by funds from the National Science Foundation (NSF: EF-2126387, IIS-2239114, and CNS-1338099), National Institutes of Health (NIH P01-AI152999), and a Centers for Disease Control (CDC) contract 75D30121C11180. KC was also supported by the Ken Kennedy Institute Computational Science & Engineering Recruiting Fellowship. BK was also supported by the National Library of Medicine Training Program in Biomedical Informatics and Data Science (T15LM007093). EKM was supported by the State of Maryland.

## A Supplemental materials

### A.1 Details of Experimental Study

#### A.1.1 Benchmarking commands

Lemur was run as:

~~~
python lemur.py -d path-to-db --tax-path path-to-taxonomy
                -t 8 --minimap2-AS -r species r
                -i path-to-input.fastq --nof -o outdir
~~~

Magnet was run as:

~~~
python magnet.py -c lemur report file -i input fastq file -o output path -m ont
~~~

Kraken 2 was run as:

~~~
kraken2 --db kraken2-db/k2 fungistd --threads 8
        --output path-to-out.txt --report
        path-to-out.report.txt path-to-input.fastq
~~~

Melon was run as:

~~~
melon -d path-to-db -t 8 -o outdir path-to-input.fastq
~~~

Centrifuger was run as:

~~~
centrifuger -u path-to-input.fastq
            -x centrifuger-db/cfr hpv+gbsarscov2
            -t 8 -o outdir
~~~

#### A.1.2 pbsim3 settings

The following command was used for generating reads with pbsim3:

~~~
pbsim3 --seed 42 --strategy wgs --depth 50.0 --method qshmm
       --qshmm pbsim3/data/QSHMM-ONT.model --length-mean 4050
       --length-sd 1000 --difference-ratio 39:24:36 --accuracy-mean 0.95
       --id-prefix {accession}
        --prefix ../Data/Simulated/Base-reads-MAGs/{accession}/{accession}
       --genome ../Data/Genomes/ncbi datasets-MAGs/data/{*accession*}/*.fna
~~~

#### A.1.3 NCBI Taxonomy

To convert scientific names to NCBI taxonomy identifiers and backwards, the Environement for Tree Exploration (ETE) toolkit was used (Huerta-Cepas et al. 2016). The corresponding taxa.sqlite NCBI taxdump was obtained on 01/18/2024 and has been fixed for all evaluations.

### A.2 Details of Experimental Results

#### A.2.1 Additional Analysis of Human Gut Microbes

